# Stress resilience is associated with transcriptional remodeling in the VTA

**DOI:** 10.1101/2025.11.13.687606

**Authors:** Adelaide R. Minerva, Brenna McMannon, Rixing Lin, Anna Zhukovskaya, Ilana B. Witten, Catherine Jensen Peña

## Abstract

Individual responses to chronic stress vary, with some individuals remaining resilient while others exhibit susceptibility. The ventral tegmental area (VTA), a region involved in reward learning, and the lateral habenula (LHb), a region involved in aversive learning, have been implicated in the pathophysiology of stress-related mood disorders. Here, we seek to understand the molecular adaptations in these regions at the level of single cells that mediate susceptibility versus resilience. In particular, it remains unclear whether, at the level of gene expression, different cell types within different brain regions mediate stress susceptibility versus resilience, or if these phenotypes are mediated by distinct trajectories within the same cell types. To address this gap, we performed single-nucleus RNA-sequencing of LHb and VTA of mice subjected to chronic social defeat stress. While we found minimal gene expression changes in the LHb after stress, the VTA exhibited widespread, cell type-specific transcriptional remodeling in resilient individuals and few gene expression changes in susceptible individuals. Across VTA cell types, resilience was associated with the coordinated upregulation of genes involved in intercellular signaling and neural communication, with maintenance of receptor-ligand interaction strength in resilience that was not present in susceptibility. Within VTA neurons, gene expression changes were most prominent in glutamatergic and dopaminergic clusters. Multivariate analyses of dopamine and glutamate subclusters showed that resilient neurons diverged more from control than susceptible neurons, but along a similar trajectory, supporting a model in which resilience reflects greater stress-related adaptations in these cell types. Together, these findings highlight the VTA as a key site of molecular plasticity in stress resilience and therefore a potential therapeutic target.

## Introduction

Mood disorders are among the most pervasive and disruptive mental health disorders worldwide^1,2^. These illnesses are often triggered or exacerbated by chronic stress^3–7^. However, not all individuals who experience stress develop psychiatric symptoms^8^, and therefore an important scientific and public health goal is to identify the biological mechanisms that promote susceptibility or confer resilience.

Chronic social defeat stress (CSDS) is a well-validated paradigm that captures individual differences in stress susceptibility in rodents^9–13^. In the CSDS model, a subset of animals exhibit social avoidance and anxiety-like behavior following repeated social defeat and are considered susceptible, while the remainder are characterized by a lack of overt behavioral changes compared to control and are considered resilient.

Multiple brain regions likely mediate the heterogeneous effects of chronic stress. The lateral habenula (LHb) is an area that drives aversive learning^14–18^, is implicated in the pathophysiology of depression^19–29^, and shows stronger activity in susceptible animals during and following CSDS^30^. In contrast, the ventral tegmental area (VTA), a key node in the brain’s reward circuitry, is engaged in resilience^31–33^. During CSDS, resilient mice have higher dopamine (VTADA) activity during resilience-associated behavior, and stimulating VTADA neurons can increase the chance of resilience^32^.

These changes underlying susceptibility or resilience could arise from shifts in gene regulation. Here, we seek to distinguish between multiple hypotheses regarding the structure of such shifts. The behavioral change in susceptible mice could be accompanied by underlying gene expression changes that support strong aversive learning (such as in the LHb), while resistance to behavioral change in resilient mice may reflect only minimal changes along the same molecular trajectory (Figure 1A, Hypothesis 1). Alternatively, resilience may recruit more pronounced adaptations, but again in the same direction as susceptibility, (such as in the VTA) to counteract the effects of stress and maintain behavioral stability (Figure 1A, Hypothesis 2). Additionally, within each region, susceptibility and resilience might represent divergent molecular trajectories away from control in different directions (Figure 1A, Hypothesis 3). Under this hypothesis, the two phenotypes are not simply defined by the degree of transcriptional change in a given direction, but by fundamentally different gene expression programs.

**Figure 1.**
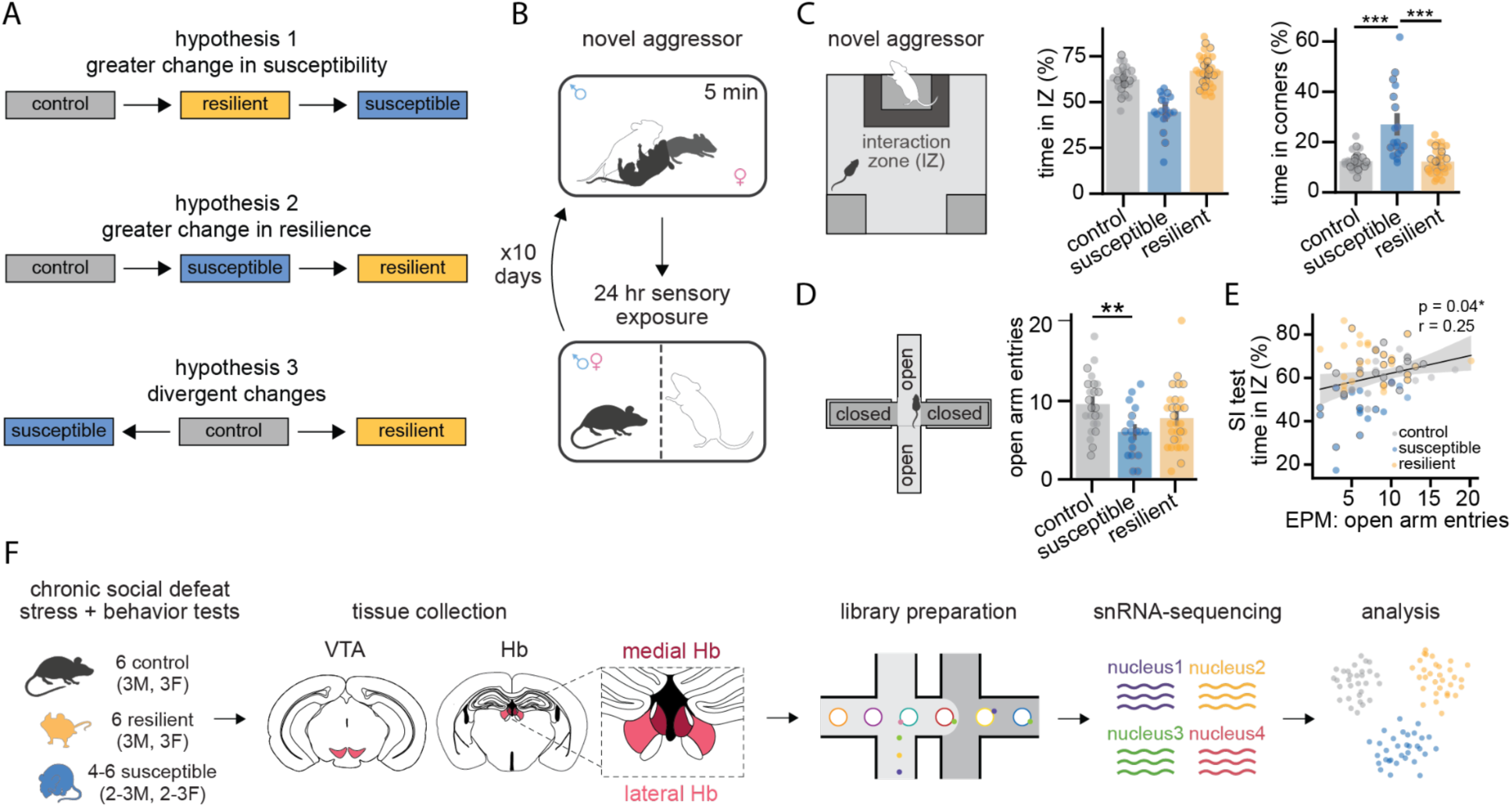
snRNA-sequencing in the VTA and LHb after chronic social defeat stress A. Hypotheses for how resilience and susceptibility may be instantiated in the brain at a transcription level. B. Schematic of chronic social defeat stress paradigm. C. During the social interaction test, time spent in the interaction zone (left) and corners (right) of the arena (1-way ANOVA for time in corners with behavior outcome as factors, F(2,69)=21.69, p=4.9e-08; Tukey HSD for control vs resilient p=1.0, control vs susceptible p=0.0, resilient vs susceptible p=0.0). Black outlines indicate which mice were used for sequencing. D. In the elevated plus maze, number of entries into the open arms (1-way ANOVA for open arm entries with behavior outcome as factors, F(2,69)=4.89, p=0.01; Tukey HSD for control vs resilient p=0.175, control vs susceptible p=0.008, resilient vs susceptible p=0.255). Black outlines indicate which mice were used for sequencing. E. Relationship between time in the interaction zone during the SI test and number of entries into the open arms of the EPM (Pearson correlation, r=0.247, p=0.036). Black outlines indicate which mice were used for sequencing. F. Schematic of tissue collection, library preparation, and snRNA-seq. See Table S1 for statistics. ∗p < 0.05, ∗∗p < 0.01, ∗∗∗p < 0.001. Unless specified, data plotted as mean ± SEM.

To test these hypotheses, we characterized transcriptomic adaptations in the LHb and VTA following chronic social stress. Cells in both regions are molecularly^34–43^ and functionally diverse^15,17,21,40,44–82^ and therefore it is likely that subpopulations may exhibit unique adaptations. Most prior work on gene expression changes underlying stress have used bulk tissue approaches^10,83–85^ and thus would be unable to test these hypotheses directly. Therefore, unbiased study of all cell types in the VTA is necessary for a comprehensive understanding of stress-induced, phenotype-specific gene expression changes. To address this, we used single-nucleus RNA-sequencing (snRNA-seq) to profile transcriptional changes across cell types in both regions of mice resilient and susceptible to CSDS. More specifically, we test the idea that different regions and cell types primarily underlie stress susceptibility (Figure 1A, Hypothesis 1), primarily underlie stress resilience (Figure 1A, Hypothesis 2), or contribute to both processes by undergoing different types of changes (Figure 1A, Hypothesis 3).

Surprisingly, we found few gene expression changes in the LHb following stress. In contrast, in the VTA we found that (1) resilience exhibits more changes in gene expression than susceptibility, (2) these changes are most dramatic in glutamatergic and dopaminergic VTA neurons, and (3) the direction of transcriptional remodeling in VTA DA and glutamate neurons is similar in susceptibility and resilience. These findings support a model in which the VTA contributes to resilience through active, cell type-specific molecular adaptations in response to stress which serve to maintain behavioral stability in the face of adversity.

## Results

### CSDS produces susceptible and resilient groups

To assess resilience versus susceptibility to stress, we subjected mice to CSDS. Mice underwent ten consecutive days of defeat from novel aggressors (Figure 1B) followed by 3-8 days of behavioral testing (Figure S1A). The social interaction test (Figure 1C, S1B-E) was used to determine stress susceptibility and mice showed a spectrum of social interaction times with a novel aggressor following CSDS. In concordance with prior literature^30,32^, stressed mice were defined as susceptible if the time they spent in the interaction zone was less than one standard deviation below that of unstressed controls of the same sex; otherwise, they were considered resilient (Figure 1C, S1C). Sex-specific thresholds were used because female mice were overall more social than males (two-way ANOVA with sex and stress as factors; main effect of sex, F=5.32, P=0.02). Unstressed male and female mice spent 60.23±8.3% (mean±s.d.) and 64.29±6.12% of time in the interaction zone near the novel aggressor, respectively. Using this measure, similar proportions of male and female mice were assigned to each group (Figure S1E). Susceptibility by this measure corresponded with anxiety-like behavior in non-social settings, such as the elevated plus maze (EPM; Figure 1D-E, S1J-M, S1O-P) and open field test (OFT; Figure S1F-I, S1N). However, it should be noted that behavior in these additional tests did not perfectly differentiate resilient from susceptible mice, likely due to behavioral complexity and individual variation in rodent models of stress.

### Transcriptional atlases of the mouse VTA and LHb using single-nucleus RNA-sequencing

To examine how stress affects both the VTA and LHb, we generated snRNA-seq atlases of transcriptional diversity in both regions from ∼10 week old male and female C57BL/6J mice. Tissue was collected from control, susceptible, and resilient mice 10-15 minutes after a final social defeat re-exposure (or conspecific exposure for controls; Figure 1F). We then isolated nuclei to generate snRNA-seq libraries and performed sequencing. Following quality control and preprocessing, we retained a total of 162,413 VTA nuclei across 16 samples and 228,187 habenula nuclei across 18 samples.

VTA nuclei clustered into 3 neuronal and 6 non-neuronal populations (Figure 2A, S2A-I; median unique molecular identifiers [UMIs]/nucleus=1,223; median genes/nucleus=821; Figure S2J). These populations, including well characterized glial cells (Figure S2), were recovered in proportions similar to those from previous literature^39^ (Figure 2B) and were similarly represented across behavioral groups (Figure S3A-C).

**Figure 2.**
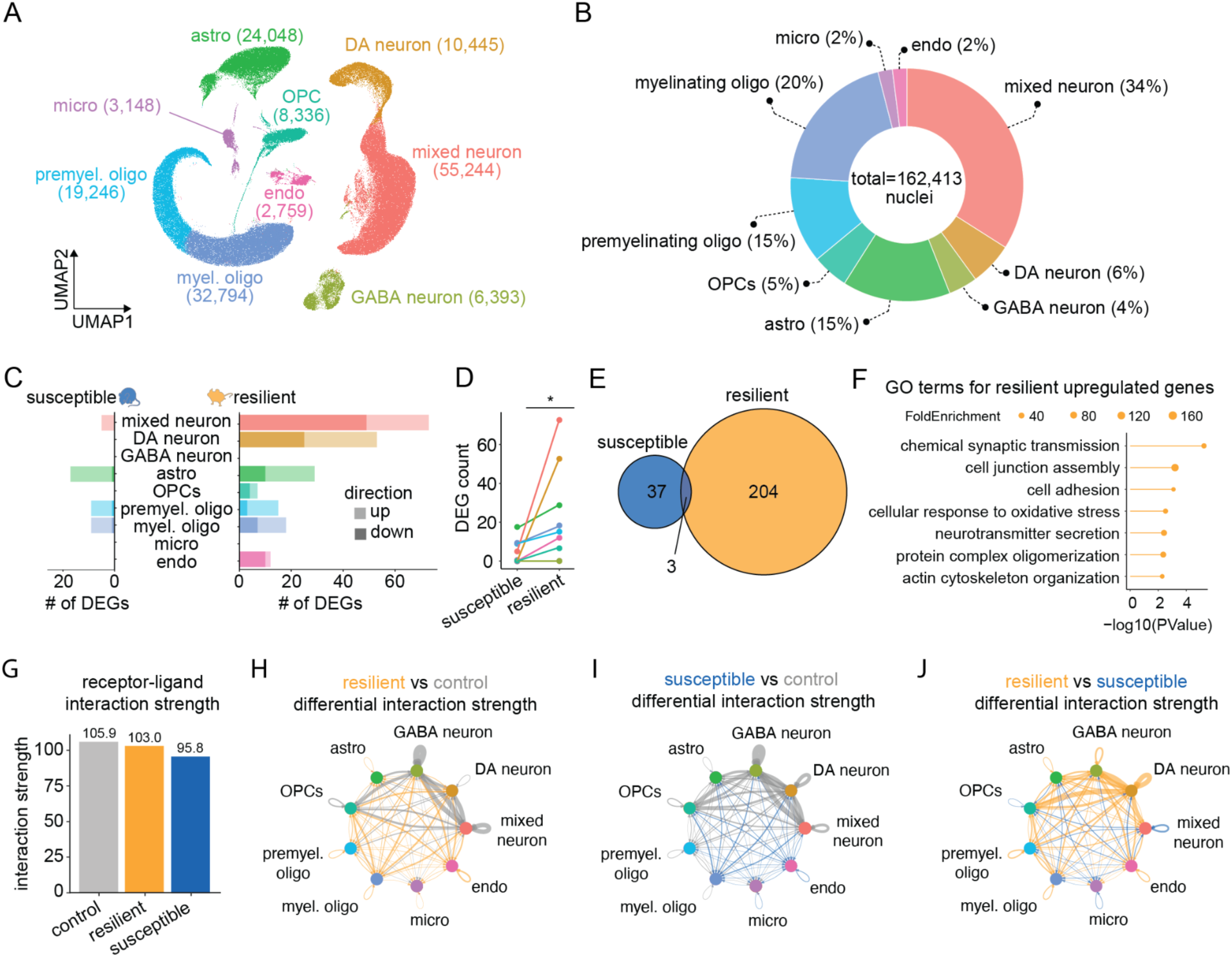
Across VTA cell types, resilient mice have greater gene expression changes and maintain receptor-ligand interaction strength. A. UMAP of all VTA nuclei with points colored by cell type classification. B. Proportion of VTA nuclei belonging to each cluster (n=162,413 total nuclei). C. Number of DEGs compared to control in susceptible (left; mixed neuron=5, DA neuron=0, GABA neuron=0, astro=17, micro=0, endo=0, OPC=0, premyelinating oligo=9, myelinating oligo=9) and resilient (right; mixed neuron=73, DA neuron=53, GABA neuron=0, astro=29, micro=0, endo=12, OPC=7, premyelinating oligo=15, myelinating oligo=18). D. DEGs counts in susceptible versus resilient groups compared to control, by cell type (Wilcoxon rank sum test with continuity correction, W=18, P=0.045; ∗p < 0.05). See Table S1 for statistics. E. Venn diagram of susceptible and resilient DEGs. F. Top GO terms associated with genes upregulated in resilience. G. Overall receptor-ligand interaction strength between all cell types. H-J. Circle plots showing differential receptor-ligand interaction strength between all cell types each pair of behavioral outcomes. Resilient nuclei maintain interaction strength similar to that of control (H; similar overall gray and gold edge width); susceptible nuclei have reduced interaction strength compared to control (I; greater overall gray than blue edge width); resilient nuclei have higher interaction strength than susceptible nuclei (J; greater overall gold than blue edge width).

Consistent with prior reports^86^, our dataset showed that the overall cell type composition of the VTA is not significantly different between the sexes (Figure S4A-C). However, the VTA shows sex-dependent variation in response to stress^55,87^ and drugs of abuse^88–91^, and receives input from brain regions that show substantial sex differences^92^. Therefore, we asked whether VTA cell types show sex-specific transcriptional signatures. Within each cluster, we pseudobulked data and performed pairwise comparisons of gene expression between males and females to reveal differentially expressed genes (DEGs) between the sexes (Figure S4D). The most dramatic sex differences were in known sex-linked genes (e.g. *Tsix* and *Uty,* Figure S4E-F). We also identified autosomal genes with strong sex bias across all cell types (e.g. *Fkbp5*, which plays a key role in stress-related disorders^93^ and has sex-dependent effects on stress susceptibility^94^; Figure S4G) as well as those that exhibited sex-specific expression in certain cell types (e.g. *Ube3a*, Figure S4H). These differences may be associated with sex-dependent responses of the VTA in different physiological or pathological contexts.

Habenula nuclei also clustered into distinct neuronal and non-neuronal populations (Figure S5A, left). We used known marker genes^35,36,43^ to distinguish neurons from medial and lateral Hb (MHb and LHb, respectively) that were distinct from extraneously captured neurons from surrounding regions (Figure S5A). Since we could not be confident about which subregion non-neuronal cell types came from due to a lack of robust markers, we excluded them from further analysis. To examine neuronal heterogeneity within both MHb and LHb and ensure consistency with prior datasets^35,36,43^, we subclustered Hb neurons (Figure S5A, right). Subclustering revealed 7 transcriptionally distinct populations (4 LHb, 2 MHb, and one that could not be confidently assigned to either subregion) consisting of 108,270 total nuclei and 23,318 genes (UMIs/nucleus=2732; median genes/nucleus=1557; Figure S5F). All subclusters expressed high levels of *Slc17a6* (Vglut2; Figure S5B,E) and low levels of GABAergic markers (Figure S5E), with LHb and MHb subclusters marked by expression of canonical markers *Pcdh10* and *Tac2*, respectively (Figure S5C-E). We also recapitulated discrete MHb subtypes expressing *Chat* or *Calb2* (Figure S5E). Subpopulations were equally represented across sexes (Figure S5G) and behavioral groups (Figure S5H).

### LHb does not show strong patterns of gene expression change with stress

Given significant research implicating hyperactivity of the LHb in aversive processing^14–18^, depression^17,20–22,61,64,95^, and stress susceptibility^23,30,96,97^, we asked whether stress there were significant gene expression changes in this region following stress. One study found divergent stress-associated gene expression in LHb, which hints at Hypothesis 3^23^. Our high resolution transcriptional atlas allowed us to also probe changes in MHb in addition to LHb neurons. Within LHb and MHb neurons separately, we performed pairwise comparisons of pseudobulked gene expression between control versus susceptible, control versus resilient, and resilient versus susceptible, including sex as a covariate. Unexpectedly, both regions showed few gene expression differences between conditions (adjusted pvalue<0.1; Figure S5I-N). We then calculated DEGs between LHb and MHb control samples and detected 5164 DEGs, indicating that our samples were sufficiently powered to detect large differences in habenula. Despite a lack of DEGs, LHb neurons of susceptible mice tended to be more active than those of resilient mice in response to a final aggressor exposure (measured by IEG expression; Figure S5O).

### Resilient mice show greater gene expression changes than susceptible across all VTA cell types while maintaining receptor-ligand interaction strength

We then wondered if the VTA, a region associated with reward rather than aversion processing, may show greater transcriptional changes following stress. We first focused on both neuronal and non-neuronal clusters in order to understand how stress affects the entire VTA network. Within each cluster, we performed pairwise comparisons of pseudobulked gene expression between control versus susceptible and control versus resilient, including sex as a covariate. Resilience was consistently associated with a greater number of DEGs in all cell types where DEGs were detected (Figure 2C-E). To ensure that the higher number of DEGs in resilience was not due to one extra resilient sample of each sex (6 resilient vs 4 susceptible), we systematically removed every combination of one male and one female resilient sample and re-calculated DEGs. Across these held out combinations, the number of resilient DEGs remained significantly greater than susceptible DEGs (Figure S6).

Resilient DEGs in non-neuronal cell types (Figure S7A-F) included several genes with putative roles in supporting network stability and function. For example, *Grm5* (metabotropic glutamate receptor 5), which plays a role in modulating synaptic plasticity and glutamate homeostasis, was upregulated in astrocytes (Figure S7A,G). This finding is consistent with work implicating astrocytic glutamate regulation in stress response and psychiatric disease^98–100^. *Ptgfrn* is involved in cell adhesion and was upregulated in OPCs (Figure S7B,H). The glucocorticoid-responsive kinase *Sgk1*, which was upregulated in myelinating oligodendrocytes (Figure S7D,I), has been implicated in processes important for maintaining circuit integrity such as myelin plasticity and stress hormone signaling^101–105^. *Bmp6* was altered in endothelial nuclei (Figure S7F,J) and is a regulator of blood-brain barrier function, potentially suggesting vascular adaptations in resilience that promote behavioral stability.

We performed gene ontology (GO) analysis on resilient DEGs to more systematically investigate the biological processes most altered in resilience. Across all clusters, genes upregulated in resilient animals were primarily associated with processes related to neural signaling and intercellular communication (Figure 2F), including chemical synaptic transmission (GO term: 0007268), cell junction assembly (GO term: 0034329), and cell adhesion (GO term: 0007155). The coordinated upregulation of genes involved in the molecular interfaces between cells and intercellular signaling in resilience may reflect the promotion of stable and efficient neural communication in order to encourage an adaptive stress response.

In order to further probe cellular communication, we used CellChat^106,107^ to compare inferred cell-cell interactions across the VTA network between each behavioral group (control, susceptible, resilient). CellChat revealed a decrease in overall interaction strength in susceptibility compared to control (Figure 2G,I), while resilience maintained interaction strength similar to that of control (Figure 2G-H). Comparing resilient and susceptibility directly highlighted stronger interactions in resilience (Figure 2J), supporting the hypothesis that the gene expression changes seen in resilience allow the VTA to maintain this interaction strength.

### Resilience is associated with greater gene expression changes in VTA neurons

We found the greatest number of DEGs and the strongest cell-cell interaction strength within neuronal clusters and, given the diversity of neuronal subtypes in the VTA, we wanted to probe these effects with finer resolution. Subclustering of VTA neurons revealed a large dopaminergic (DA) cluster as well as glutamatergic, mixed, and multiple GABAergic populations (Figure 3A-E).

**Figure 3.**
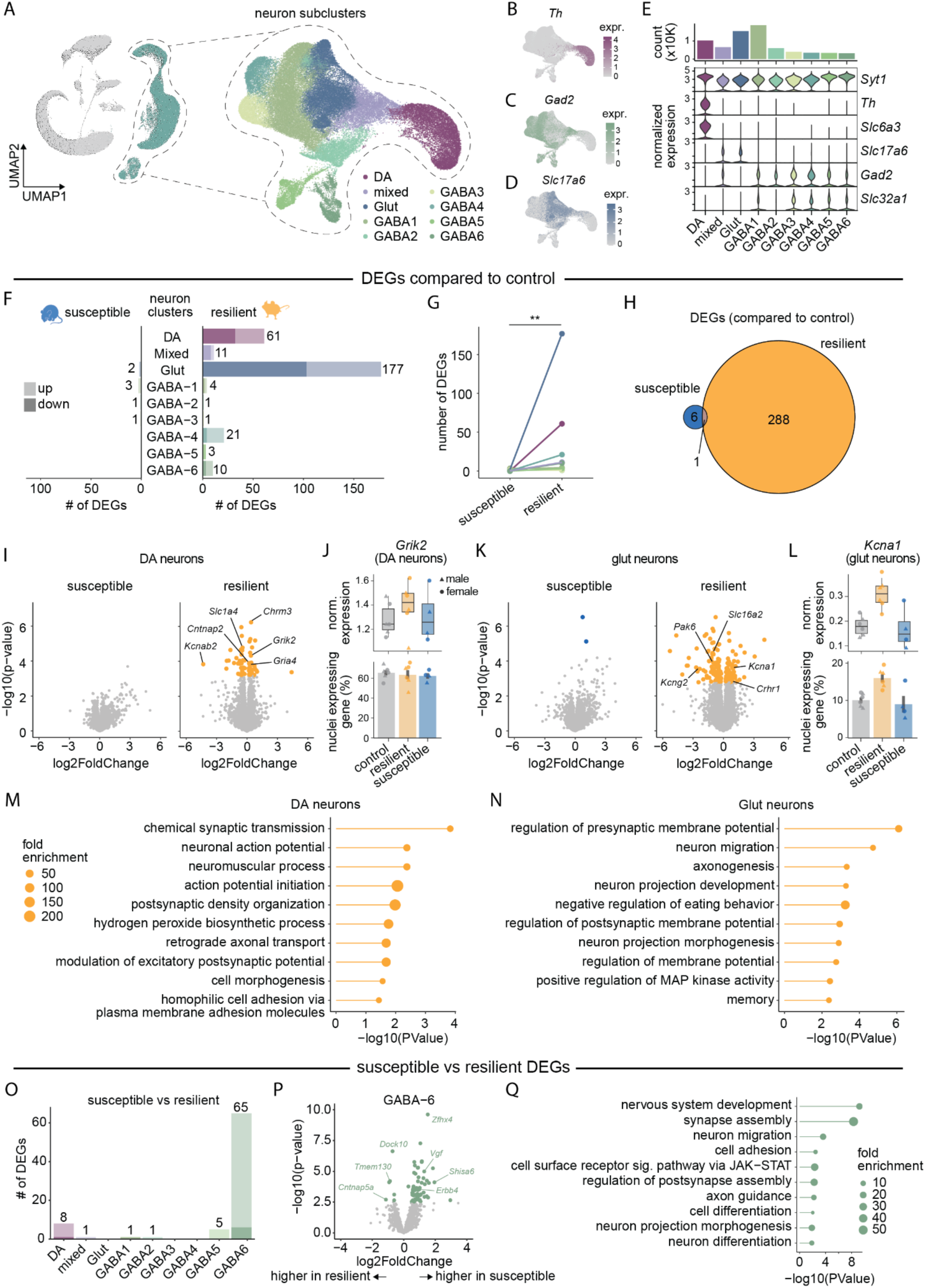
Resilience is associated with greater transcriptional changes in VTA DA and glutamate neurons, while Sst-expressing GABA neurons show differences between susceptibility and resilience. A. UMAP of all VTA nuclei with neurons highlighted (left) and neuronal subclusters (right) with points colored by cell type classification. B. Feature plot showing expression of Th across VTA neuron UMAP space. C. Same as B, but for Gad2. D. Same as B-C, but for Slc17a6 (Vglut2). E. Number of nuclei per VTA neuronal subcluster (top) and expression of select marker genes across VTA neuronal subclusters (bottom). F. Number of DEGs compared to control in susceptible (left; DA=0, Mixed=0, Glut=2, GABA1=3, GABA2=1, GABA3=1, GABA4=0, GABA5=0, GABA6=0) and resilient (right; DA=61, Mixed=11, Glut=177, GABA1=4, GABA2=1, GABA3=1, GABA4=21, GABA5=3, GABA6=10) neurons. G. Summary of DEGs counts in susceptible versus control and resilient versus control, by cell type (Wilcoxon rank sum test with continuity correction, W=6.5, P=0.003; ∗∗p < 0.01). See Table S1 for statistics. H. Venn diagram of susceptible and resilient DEGs. I. Volcano plots showing susceptible DEGs (left) and resilient DEGs (right) in DA neurons. Nominal p-values are plotted, but points are colored based on adjusted p-values. Adjusted p-value (Bonferroni post hoc correction) cutoff is 0.1. J. Expression of the DA DEG Grik2 across behavioral groups. Top: each point represents the normalized expression of Grik2 across all DA neurons in each sample. Differential expression was assessed using DESeq2 on pseudobulked counts (log2FC=0.38, Wald statistic=4.07, p.adjusted=0.03). Bottom: each point represents the proportion of DA nuclei in each sample that express Grik2. K. Same as I, but for glutamate neurons. L. Same as J, but for the glutamate DEG Kcna1 (log2FC=0.94, Wald statistic=3.64, p.adjusted=0.04). M. Top GO terms associated with genes altered in DA neurons of resilient mice compared to control. N. Top GO terms associated with genes altered in glutamate neurons of resilient mice compared to control. O. Number of DEGs between susceptible and resilient (DA=8, Mixed=1, Glut=0, GABA1=1, GABA2=1, GABA3=0, GABA4=0, GABA5=5, GABA6=65) neurons. P. Volcano plot showing DEGs between susceptible and resilient in GABA6 neurons. Nominal p-values are plotted, but points are colored based on adjusted p-values. Adjusted p-value (Bonferroni post hoc correction) cutoff is 0.1. Top GO terms associated with genes differentially expressed in GABA6 neurons of resilient versus susceptible mice.

The six GABAergic clusters, marked by high levels of *Gad2* and *Slc32a1* (Figure 3C,E, S8A-H), reflected the transcriptomic diversity of VTA GABA neurons. These neurons modulate local DA activity^65,66,108,109^, send long-range projections^75,110–116^, play diverse roles in processing reward and aversion^53,65,66,70–72,74,76^, and exhibit transcriptomic heterogeneity^117–120^. Two clusters (GABA3 and GABA4) expressed high levels of *Htr2c* (Figure S8B,H) and likely represent populations previously identified in the VTA^39,121^. GABA5 and GABA6 expressed canonical interneuron markers such as *Sst*, *Kit*, *Nos1*, *Oprm1*, *Vip,* and *Sox6*^122^ (Figure S8F-H). While these genes typically mark non-projecting GABA neurons in other brain regions, there is still debate surrounding whether canonical interneurons exist within the VTA and whether GABAergic VTA projection neurons also have local collaterals that can modulate the activity of neighboring neurons^123^.

To further characterize these neuronal subtypes, we mapped our neuronal nuclei to the Allen Brain Cell (ABC) Atlas using the Allen Institute’s *mapmycells* function (see Methods). The mapping results are summarized in Table S6, which lists the top ABC cell classes accounting for up to 90% of the mapping for each of our assigned clusters. The majority of our clusters corresponded to midbrain neurons of the same neurotransmitter type, consistent with accurate anatomical and molecular annotation. Clusters GABA3 and GABA5 showed predominant mapping to pons GABA neurons, which may reflect minor dissection variability or shared transcriptional signatures among inhibitory neurons across neighboring regions.

To assess cell type-specific transcriptional changes related to stress response, we performed pairwise pseudobulked comparisons of gene expression between control versus susceptible, control versus resilient, and susceptible versus resilient within each VTA neuronal subcluster. Compared to control, resilience was associated with greater or equal numbers of DEGs than susceptibility in all subclusters (Figure 3F-H).

VTA DA and glutamate neurons had the greatest number of DEGs (Figure 3F). DA DEGs included potassium channels such as *Kcnab2*, cell adhesion protein *Cntnap2* that is known to cluster around potassium channels and play a crucial role in synaptic connections, amino acid transporter *Slc1a4*, and both acetylcholine and glutamate receptors (*Chrm3*, *Grik2*, *Gria4*; Figure 3I-J). Glutamatergic DEGs included potassium channels (*Kcna1*, *Kcng2*), *Pak6* (a kinase implicated in synaptic plasticity and neurotransmission), corticotropin releasing hormone receptor *Crhr1*, and thyroid hormone transporter *Slc16a2* (Figure 3K-L).

Given that VTADA neurons are more heterogeneous than previously thought, we also looked for DA subcluster-specific DEGs. Subclustering of VTADA neurons (Figure S9A) revealed similar subpopulations to those found in previously collected sc- and snRNA-seq datasets generated from mouse and rat midbrain^37–39,124^. We observed ubiquitous expression of canonical DA neuron markers (e.g. *Th, Slc6a3, Ddc, Slc18a2*) as well as additional VTADA subpopulations previously identified to be anatomically and functionally-distinct (Figure S9B-H). While we saw the same pattern of greater change in resilience than susceptibility, we surprisingly found fewer DEGs overall among VTADA subclusters than in the higher level DA cluster (Figure S9I). Subcluster DA-2 had the highest number of DEGs (Figure S9I) and may be driving the resilience-associated changes in DA neurons, but the overall low number of DEGs at the subcluster level suggests that transcriptional changes associated with resilience may be distributed across dopaminergic subtypes rather than concentrated within specific ones. Due to the low number of DA subcluster DEGs, we conducted downstream analyses at the higher DA cluster level, but future studies may probe the role of individual DA subclusters in responding to stress.

We performed GO analysis on resilient DEGs in DA and glutamate neurons (Figure 3M-N). In resilient animals, DA neuron DEGs were associated with synaptic transmission (GO terms: 0007268, 0098815, 0097106), neuronal excitability (GO terms: 0019228, 0099610), and structural connectivity (GO terms: 0008090, 0000902, 0007157), suggesting adaptations that may fine-tune how these neurons communicate in real-time. In contrast, glutamate neuron DEGs were associated with developmental and structural plasticity processes such as neuron migration (GO term: 0001764) and projection growth (GO terms: 0031175, 0048812) as well as MAP kinase signaling (GO term: 0043410). These terms suggest long-term remodeling that might support behavioral flexibility and learning associated with resilience.

The one population that exhibited a large number of DEGs between susceptible and resilient animals was a specific population of GABAergic VTA neurons (GABA6; Figure 3O-P). This cluster was characterized by high expression of *Sst, Oprm1, Kit, Nos1,* and other peptidergic and synaptic regulatory genes (Figure S8G-H). According to our ABC mapping, GABA6 most closely corresponded to ABC midbrain (MB) GABA neurons (Table S6). This mapping reinforces that the observed transcriptional differences between resilient and susceptible animals are intrinsic to a midbrain GABAergic population rather than driven by contamination from neighboring regions. GO analysis of susceptible versus resilient GABA6 DEGs highlighted differences in neuronal development, synapse assembly, and signaling pathways, suggesting altered circuit connectivity and plasticity between the two phenotypes in this population (Figure 3Q).

### Linear discriminant analysis reveals greater molecular divergence in resilience than susceptibility compared to control

In VTA DA and glutamate neurons, we observed more DEGs between resilient and control than between susceptible and control. These findings challenge the hypothesis that susceptibility represents a greater molecular shift from the control state than resilience (Figure 1A, Hypothesis 1). In addition, the small number of DEGs between susceptible and resilient in these neuronal clusters challenge the idea that susceptibility and resilience diverge in distinct directions away from control (Figure 1A, Hypothesis 3). Instead, our DEG analysis supports the idea that resilience represents a greater shift in expression from control in these cell types (Figure 1A, Hypothesis 2). Alternatively, one population of GABAergic VTA neurons characterized by high expression of peptidergic and synaptic regulatory genes (GABA6) had few DEGs in either resilient or susceptible compared to control but showed large differences between resilient and susceptible directly (Figure 3O). This pattern suggests that susceptibility and resilience diverge more from each other than from control (Figure 1A, Hypothesis 3).

To investigate the overall transcriptional structure between behavioral groups and test these hypotheses more directly, we applied linear discriminant analysis (LDA) to each of these three clusters (DA, glutamate, GABA6). This approach allowed us to identify the axes that best distinguish all three behavioral groups (Figure 4A). In both DA and glutamate neurons, group centroids laid approximately along a linear trajectory (Figure 4B, D), exhibiting a graded shift in transcriptional state from control to susceptible to resilient that supports Hypothesis 2. In DA neurons in particular, when nuclei from susceptible mice were misclassified, they were more often labeled as control than resilient (Figure 4C), reinforcing their transcriptional proximity to control. This also occurred to a lesser extent among glutamate neurons (Figure 4E). These results suggest that a one-dimensional axis could effectively capture the main group differences in these subclusters and that, in these cell types, gene expression in susceptible mice is most similar to control.

**Figure 4.**
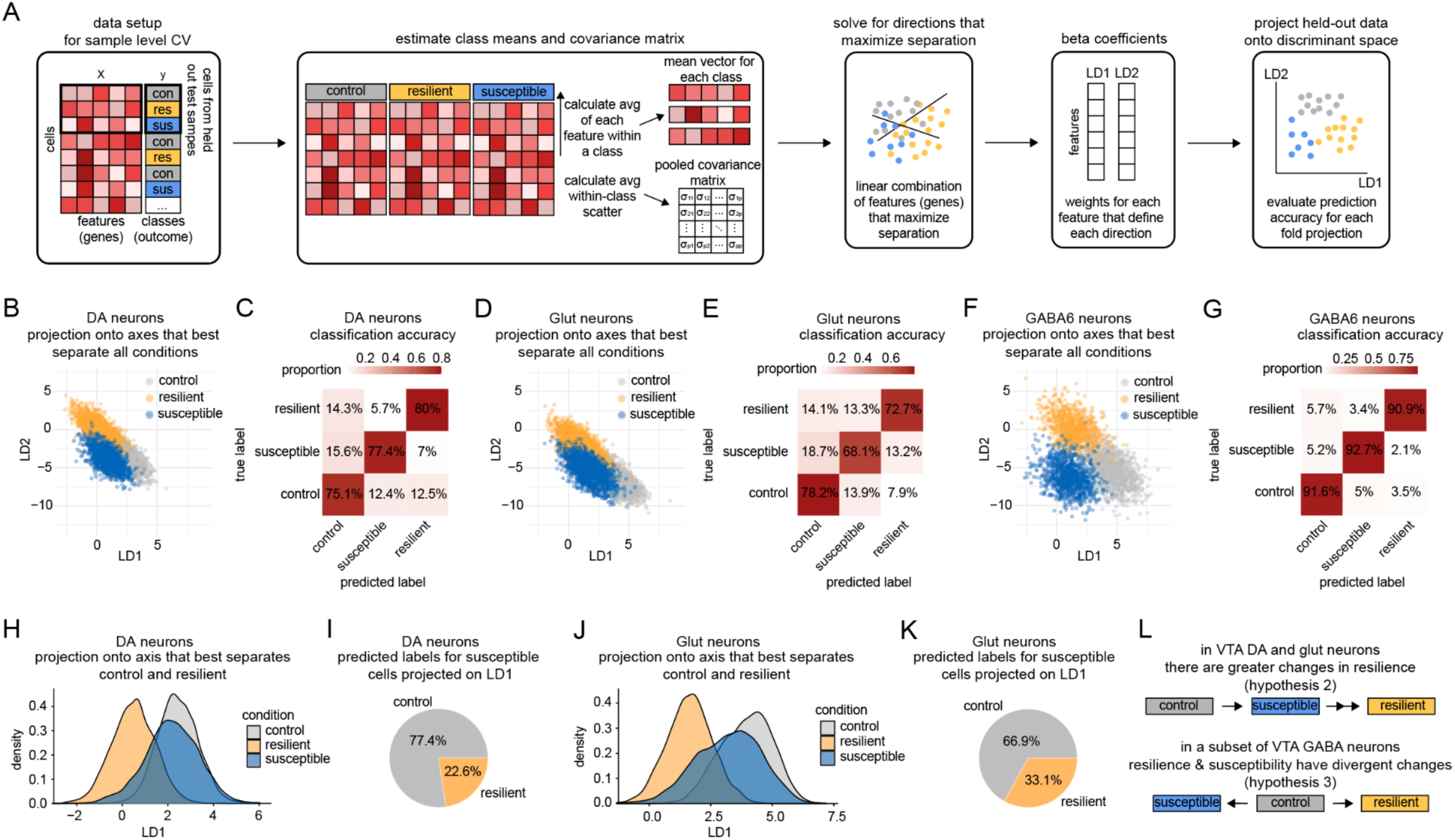
LDA identifies cell type-specific transcriptional trajectories in resilient and susceptible VTA neurons. A. Schematic of the cross-validation (CV) workflow for 2D LDA. B. 2D LDA projections of held-out nuclei from DA neurons, colored by true group labels. C. Confusion matrix showing classification accuracy of held-out DA nuclei, aggregated across all folds. D. Same as B, but for glutamate neurons. E. Same as C, but for glutamate neurons. F. Same as B, but for GABA6 neurons. G. Same as C, but for GABA6 neurons. H. 1D LDA projection of DA nuclei onto LD1 trained only and control and resilient. I. Pie chart summarizing predicted classifications for susceptible nuclei projected on the control-resilient LD1 axis. J. Same as F, but for glutamate nuclei K. Same as G, but for glutamate nuclei. LDA supports hypothesis 2 for how resilience and susceptibility are instantiated in VTA dopamine and glutamate neurons and hypothesis 3 for a specific population of VTA GABA neurons (GABA6).

To further test this idea, we performed one-dimensional LDA on DA and glutamate neurons using only control and resilient nuclei to derive the linear discriminant (LD1) that best separates these two groups. We then projected susceptible nuclei onto this axis to examine their relative positioning (Figure 4H, J). In both DA and glutamate neurons, susceptible nuclei indeed landed closer to control than to resilient along LD1, and were more often classified as control than resilient (Figure 4H-K). These results reinforce the idea that in these cell types, susceptibility maintains overall transcriptional structure much closer to control than resilience does (Figure 4L, Hypothesis 2).

In contrast, in GABA6 neurons all three group centroids occupied distinct regions in LDA space (Figure 4F) and were classified well above chance (Figure 4G). These results support the idea that resilience and susceptibility are instantiated by discrete gene expression structures away from control within this population (Figure 4L, Hypothesis 3).

Together, these analyses uncover cell type-specific transcriptional structures in VTA that distinguish susceptible and resilient phenotypes following chronic social stress.

## Discussion

Chronic stress is a significant trigger for depression and other mood disorders, and therefore the study of mechanisms that confer resilience to stress are of great importance for public health. Here, we demonstrated that resilience to chronic stress involves active, cell type-specific transcriptional changes in the VTA and is not simply a passive lack of susceptibility. Surprisingly, despite its well-established role in aversive signaling, the LHb exhibited minimal gene expression changes.

### Limited transcriptional changes in the LHb suggest alternative mechanisms of plasticity

While the LHb is widely implicated in aversive processing and stress-related behavior^14–30^, we observed few transcriptional changes in this region in either susceptible or resilient animals following CSDS. These unexpected findings suggest that the LHb may exert its influence through alternative mechanisms that do not involve sustained transcriptomic remodeling within neurons. One possibility is that stress induces transient gene expression changes that drive synaptic remodeling, which can maintain changes in neuronal activity with normal gene expression. A similar pattern of rapid transcriptional response to acute stress followed by long-term dampening of this response as stress becomes chronic has recently been reported in ventral hippocampus^125^. This underscores the importance of assessing multiple time points and complementing transcriptomic studies with circuit-level and physiological approaches to fully capture regional dynamics. In addition, studies have shown that stress-induced alterations in LHb astrocyte function can modulate LHb neuron firing and depressive-like behavior^57–59,126^. These effects may be reflected as astrocyte-specific transcriptional changes and warrant further investigation in future studies.

### Cell type specificity of stress adaptations in the VTA

Within VTA neurons, we observed distinct gene expression patterns across dopaminergic, glutamatergic, and GABAergic subpopulations. These results reveal that susceptibility and resilience diverge in their transcriptional trajectories across neuron subtypes and highlight that individual cell types within the VTA adapt to stress heterogeneously. Specifically, within VTA DA and glutamate neurons, our analyses support the idea that resilience recruits more pronounced adaptations in a similar direction as susceptibility away from control (Figure 4L, Hypothesis 2). On the other hand, in a specific population of *Sst-*expressing GABA neurons (GABA6), our data supports the hypothesis that susceptibility and resilience represent fundamentally different gene expression programs, moving away from control in divergent directions (Figure 4L, Hypothesis 3). Together, this reinforces the importance of examining unique cell types in this region to understand the precise molecular mechanisms that differentiate susceptibility and resilience to chronic stress.

### Resilience as an active adaptive response in the VTA

The VTA showed widespread transcriptional changes across both neuronal and non-neuronal cell types, with resilience consistently associated with more DEGs than susceptibility, supporting and extending earlier bulk RNA-seq findings^10^. These changes were in genes related to synaptic transmission, structural plasticity, and cell-cell communication, suggesting that resilient animals engage regulatory programs to stabilize neural function and preserve network connectivity. These changes were largest in VTA DA and glutamate neurons. Notably, susceptibility involved far fewer transcriptional changes despite producing strong behavioral phenotypes, suggesting that resilience requires a more coordinated molecular response.

Further, multivariate LDA analysis of overall transcriptional state found that within both VTA DA and glutamate neurons, group centroids laid approximately along a linear trajectory from control to susceptible to resilient. This graded positioning reinforces the asymmetry we found in the number of DEGs: greater transcriptional distance between control and resilient nuclei than control and susceptible nuclei reflects more extensive transcriptional remodeling in resilience, whereas susceptibility maintains overall transcriptional structure closer to control. In contrast, the closer proximity of susceptible nuclei to control is represented by a more subtle shift in gene expression, represented by fewer DEGs. These findings support the hypothesis that within VTA DA and glutamate neurons resilience is not merely the absence of susceptibility, but is instead an active transcriptional state that extends further from control than susceptibility does and likely engages stronger regulatory programs. This view aligns with electrophysiological findings that resilient animals maintain stable VTADA neuron firing, likely through compensatory, homeostatic mechanisms that preserve neural function under stress^33^.

These findings challenge the prevailing view that to ameliorate the changes in mood, behavior, and neural activity associated with stress susceptibility, we must reverse the molecular consequences of stress. By identifying resilience as an active and widespread transcriptional program in the VTA, our work provides a foundation for future efforts to target cell type-specific pathways that support adaptive responses to chronic stress. Whereas susceptibility may arise from a failure to activate sufficient protective mechanisms, resilience appears to involve the engagement of gene networks that promote neural stability and flexible behavioral coping. Similar forms of structural plasticity have been observed in other brain regions such as the hippocampus, where resilience is associated with compensatory postsynaptic remodeling in CA1^127^. Identifying upstream drivers of these cell type-specific transcriptional programs could inform new strategies to enhance natural resilience mechanisms. Such approaches may offer a compelling alternative to current treatments, which largely aim to suppress symptoms of mood disorders rather than promote resilience. This concept is supported by prior studies in other brain regions such as the nucleus accumbens, where similar efforts to harness endogenous resilience mechanisms have shown potential as therapeutic strategies for depression^85,128,129^.

### Caveats and Considerations

Although we performed snRNA-seq on 16 VTA samples from 32 mice and 18 un-pooled Hb samples, we are likely underpowered to detect small changes in expression given the need to correct for multiple comparisons across the genome ^130^. However, we performed several analyses to ensure that our findings are not simply due to small sample sizes. Iterative DEseq2 analysis of VTA data, in which we dropped every combination of male and female resilient samples, confirmed that an imbalance in sample size does not account for a greater number of resilient DEGs (Figure S6). The conclusion that resilience involves greater sustained transcriptional change compared to susceptibility is further supported by multivariate LDA modeling. Although we did not identify stress-associated DEGs in the Hb, comparison of gene expression between LHb and MHb control samples identified thousands of DEGs, indicating the current sample sizes are sufficient to detect the most robust differences. Nevertheless, given sample size considerations, it would be naïve to conclude that stress does not result in any gene expression differences in the VTA of susceptible mice or in the Hb of either group; rather, we can only conclude that the most robust changes are detectable among resilient mice.

Although both males and females were used and sex was specifically considered in our study design and analysis, additional considerations remain. To examine both sexes, we used chronic nondiscriminatory social defeat stress (CNSDS)^11,131^, a modified version of CSDS in which a male and a female mouse are placed together into the homecage of a novel aggressor to induce nondiscriminatory attacks, including towards the female. Although male mice tend to be attacked more than females in this paradigm, and therefore the severity of the stress may not be the same between the sexes, this approach nonetheless allowed us to use the same stressor to generate resilient and susceptible phenotypes from both sexes, which was key to the goals of this study. We controlled for the effects of sex by using sex as a factor in all DEseq2 models, but this study was not powered to examine sex-specific stress effects. Given prior evidence of sex-dependent variation in response to stress in the VTA^55,87^

Here, transcriptional profiling was performed at a single time point following CSDS, limiting the ability to capture dynamics or transient gene expression changes over time. In particular, additional time points immediately following the first stress experience, and later time points after recovery, may resolve additional signatures of susceptibility and resilience, respectively. While comprehensive longitudinal snRNA-seq remains prohibitively expensive at the scale necessary to fully capture these dynamics, reduced time-course designs or computational trajectory inference could better resolve the temporal progression of stress-induced molecular adaptations.

Finally, the analyses here have focused on characterization of the molecular signatures of resilience and susceptibility after chronic adult social stress, and have not yet functionally established the causal role of any of the specific gene expression changes observed. Future studies utilizing gene perturbation or functional assays will be necessary to test whether identified DEGs causally influence resilience.

## Acknowledgements

We thank Annegret Falkner, Fenna Krienen, Chris Zimmerman, and Lindsay Willmore for feedback, as well as support from the rest of the Peña and Witten labs. We also thank Wei Wang and Jennifer Miller of the Princeton Genomics Core for conducting library preparation and sequencing. This research was funded by NSF GRFP DGE-2039656 (A.R.M.), Princeton University Elizabeth Procter Fellowship (A.R.M.), Princeton Neuroscience Institute Research Innovator Award (C.J.P.), New York Stem Cell Foundation (C.J.P. & I.B.W.), the Howard Hughes Medical Institute (I.B.W.), NIH DP1-MH136573 (I.B.W.).

## Contributions

A.R.M, I.B.W., and C.J.P. conceived of the project and designed the experiments. A.R.M., B.M., R.L., and A.Z. collected the data. A.R.M. analyzed the data. A.R.M., I.B.W., and C.J.P. wrote the manuscript.

## Declaration of Interests

The authors declare no competing interests.

## Data and Code Availability

Single-nucleus RNA-sequencing data have been deposited at GEO and are awaiting a GEO accession number. Code for analyzing behavior and snRNA-seq data is available on GitHub: https://github.com/aminerva/CSDS-SingleCell-VTA-Hb. Any additional information required to reanalyze the data reported in this paper is available from the Lead Contact upon request.

## Supplemental Figures

Figure S1: Post hoc behavioral tests correlate with susceptible and resilient phenotypes from the social interaction test

Figure S2. Characterization and quality control metrics from snRNA-sequencing of VTA

Figure S3. Quality control metrics from snRNA-sequencing of VTA across behavioral groups

Figure S4. Sex differences across VTA cell types

Figure S5. Gene expression in the Hb does not uniquely define resilience or susceptibility

Figure S6. Differences in resilient vs control and susceptible vs control DEG counts is not due to sample size differences

Figure S7. Example DEGs in non-neuronal cell types of the VTA

Figure S8. GABAergic neuron diversity in the VTA

Figure S9. DA neuron diversity in the VTA

## Supplemental Tables

**Table S1:** Stats for main figures

**Table S2:** Stats for supp figures

**Table S3:** DESeq2 results for all VTA cells

**Table S4:** DESeq2 results for VTA neurons

**Table S5:** DESeq2 results for Hb

**Table S6:** Results from mapping neuronal clusters to the Allen Institute’s brain atlas cell classes

## Methods

### Mice

All experiments were approved by the Princeton University Institutional Animal Care and Use Committee and were in accordance with National Institutes of Health standards. Prior to and throughout experimental assays, animals were housed under a 12H light-dark cycle. Food and water were given *ad libitum*. Experimental animals used in this study were comprised of 72 mice (36 males and 36 females) on a C57/BL6 WT background (Jackson Laboratory) between the ages of 10 and 16 weeks. Tissue from 32 of these mice was used for sequencing. Aggressor mice were Swiss Webster male retired breeders (Taconic).

### Chronic social defeat stress (CSDS)

Both male and female mice were exposed to chronic non-discriminatory social defeat stress (CNSDS)^11,131^, a modified version of chronic social defeat stress^9^. Male mice were placed in the shoebox home cage (#5 Expanded Mouse Cage 22.2cm x 30.8cm x 16.2cm, Thoren Caging Systems, Inc.) of a novel aggressor mouse for 3 minutes of free interaction, followed by an additional 5 minutes during which a female was added. While aggressor mice do not reliably attack females on their own, the inclusion of both a male and a female induces nondiscriminatory attacks from the aggressor. Males were separated from the aggressor by a perforated acrylic barrier (Tap Plastics) within the aggressor cage, while females were housed in a new cage across an acrylic barrier from a different aggressor with a handful of soiled bedding from the aggressor they were defeated by that day. These housing conditions remained overnight. 24H later, mice were placed in the cage of a new aggressor and males and females were rotated in different directions so that trios were unique each day. This continued for a total of 10 days. Unstressed control mice were pair-housed with a perforated barrier separating the two mice. They were handled and their cages rotated each day for 10 days. Following the defeat on day 10, aggressors were removed and all mice were singly housed through the remaining stages of the experiment.

### Social interaction test

To assess social avoidance behavior following the final day of CSDS, experimental mice were tested in a two-stage social interaction test during their light cycle from ∼9am-12pm. In the first 2.5 min stage, experimental mice were allowed to freely explore an arena (44 × 44 × 20 cm) containing a novel plexiglass and wire mesh enclosure (10 × 6 cm) centered against one wall of the arena. The mouse was then removed from the chamber, and a novel Swiss Webster aggressor mouse was placed within the plexiglass and mesh enclosure. The experimental mouse was then returned to the arena for an additional 2.5 min. Mouse location was tracked via Ethovision (Noldus) and used to measure time spent in the “interaction zone” (14 × 26 cm) surrounding the enclosure, “corner zones” (10 × 10 cm), and “distance traveled”. Time spent in the social interaction zone when the social target was a novel Swiss Webster aggressor was used to delineate resilient and susceptible mice. We defined susceptible as 1 standard deviation below the mean social interaction time of the unstressed control group as done in previous literature^30,32^.

### Elevated plus maze

Mice were tested in the elevated plus maze (EPM) during their light cycle from ∼9am-12pm (85-101 lux in open arms). Each mouse was placed in the center of the EPM (2 enclosed arms and 2 open arms; each arm 76cm long and 6.5cm wide). The mouse explored the maze for 7 minutes, while its centroid location was tracked via Ethovision (Noldus) and used to measure the time spent in the open arms and center of the maze.

### Open field test

Mice were tested in the open field test (OFT) during their light cycle from ∼9am-12pm (85-101 lux). Each mouse was placed into the center of an empty area (50cm x 50cm). Lamps were used to illuminate the arena on the left and right so that there was a shaded area along the left and right walls. The center was 42cm x 42cm centered at the center of the arena. The animal explored for 10 minutes, while its centroid location was tracked via Ethovision (Noldus) and used to measure the time spent in the center of the arena.

### Tissue collection and single-nuclei dissociation for sequencing

We collected VTA and habenula tissue from male and female mice in adulthood (>P60). Following 10 days of CSDS, mice underwent 3-8 days of behavioral testing (described above). On day 19, mice were re-exposed to a novel aggressor (or novel conspecific for control mice) for 5 minutes. ∼10-15 minutes after re-exposure, animals from all conditions were cervically dislocated. This time delay is in line with prior reports indicating sufficient IEG exposure at this time following a social exposure ^132^. Brains were extracted and sectioned into 1mm coronal sections on ice using a brain block. Bilateral punches of VTA and one punch encompassing both hemispheres of habenula were taken, flash frozen in Eppendorf tubes, and kept at -80C until nuclei isolation and library preparation.

Nuclei were isolated into suspensions for snRNA-seq as previously described,^133^ with minor modifications. For VTA, tissue punches from two animals of the same sex and condition were pooled together to ensure a high enough concentration of nuclei per sample for quality sequencing and were then Dounce homogenized. Pairs were selected randomly from within each sex and condition. The sample was filtered through a 40um strainer and nuclei were isolated via iodixanol gradient centrifugation. For habenula, a subset of the animals pooled for VTA samples were used for sequencing and each sample contained tissue from only one animal. Nuclei isolation continued as above, except that nuclei were pelleted by centrifugation rather than isolated via gradient. Nuclei were hand-counted using a hemocytometer on an EVOS M5000 microscope and the volume of suspension used for library preparation was adjusted in order to load a target of 20,000 nuclei and capture 10,000 nuclei per sample.

### snRNA-seq library prep and sequencing

Nuclei were captured and RNA was reverse transcribed and barcoded for library preparation using the 10X Genomics Chromium v3 (VTA) or v4 (habenula) platform according to the manufacturer’s instructions. cDNA was amplified and adapters were added for sequencing on an Illumina NovaSeq SP 100nt Lane v1.5. Library preparation and sequencing was performed by the Princeton University Genomics Core.

### snRNA-seq read mapping

The 10X Genomics package CellRanger (v6.1.1) was used to map transcripts to the mm10 mouse reference genome. In order to capture the high percentage of pre-spliced intronic mRNA present within the nucleus, the ‘–include-introns’ flag was used to map unspliced reads to corresponding genes.

## QUANTIFICATION AND STATISTICAL ANALYSIS

### Statistical analysis of behaviors

All statistical tests of behavior (SI test, EPM, OFT) were analyzed using custom Python scripts. To assess differences in each behavioral metric across groups, a one-way analysis of variance (ANOVA) was performed with behavioral metric as the dependent variable and behavior group as the independent factor. When a significant main effect was observed, post hoc comparisons were conducted using Tukey’s Honestly Significant Difference (HSD) test to determine pairwise group differences. These statistical analyses were performed using the statsmodels module. To assess correlations between behavioral metrics, a Pearson correlation analysis was performed between the two metrics. Statistical values were computed using the scipy.stats module.

### snRNA-sequencing analysis

#### snRNA-seq read mapping

The 10X Genomics package CellRanger (v6.1.1) was used to map transcripts to the mm10 mouse reference genome. In order to capture the high percentage of pre-spliced intronic mRNA present within the nucleus, the ‘--include-introns’ flag was used to map unspliced reads to corresponding genes. After quality control and initial filtering, we recovered a total of 228,187 habenula nuclei and 162,413 VTA nuclei.

#### Preprocessing

CellRanger output count matrices were further analyzed using Seurat (v4.0.0)^134,135^ in R (v4.0.3). During microfluidic encapsulation of nuclei on 10X devices, some droplets may contain more than one nucleus. During reverse transcription, transcripts from co-encapsulated nuclei are labeled with the same cell barcode, creating a ‘doublet’. To remove likely doublets in each sample, nuclei with >2500 genes were removed. In addition, nuclei with <200 genes and >5% of reads mapping to mitochondrial genes were removed. Mitochondrial and ribosomal genes were then removed in order to remove these genes as a confounding source of variation in downstream analysis.

#### Dataset integration, UMAP embedding, and clustering

VTA and Hb datasets were analyzed separately. To integrate all data from the same brain region and assign cell type identity to nuclei, each sample dataset was log normalized and the top 2000 highly variable features were identified for downstream sample integration using the *FindVariableFeatures()* function. These features were used as input to the function *SelectIntegrationFeatures()*. We then scaled and ran PCA on each dataset separately. The 16 separate VTA datasets–6 control (3 male and 3 female), 6 resilient (3 male and 3 female), and 4 susceptible (2 male and 2 female)–and 18 separate habenula datasets–6 control (3 male and 3 female), 6 resilient (3 male and 3 female), and 6 susceptible (3 male and 3 female)–were then integrated using the *FindIntegrationAnchors()* and *IntegrateData()* functions. Next, we scaled and ran PCA on both integrated datasets. The *RunUMAP()* function was used to embed the nuclei into UMAP space and the *FindNeighbors()* and *FindClusters()* functions were used to cluster nuclei based on overall gene expression similarity. Known marker genes were used to assign cell type identities to each cluster.

#### Quality control metrics

In both datasets, nuclei from male and female samples were distributed evenly across clusters (Figure S4A-C, S5G). Nuclei were similarly well distributed across samples from different behavioral groups (control, susceptible, resilient; Figure S3A-C, S5H).

#### Calculation of DEGs between sexes across all VTA cell types

To identify DEGs between males and females in each VTA cluster, we performed pseudobulk DESeq2 analysis similar to above, but with slight modifications. The design formula included behavioral condition as a covariate to adjust for stress-specific effects: `design = ∼ sex + condition.` Normalized counts were calculated and genes were filtered based on expression thresholds to improve statistical power: genes with normalized counts above the 5th percentile in at least four samples were retained. The same FDR adjusted p-value threshold was used to determine significance.

#### Identification of Hb neuronal subtypes

We further subclustered neurons in the Hb dataset to identify neuronal subtypes in both the LHb and MHb. Clusters were identified as neuronal based on expression of *Syt1* and as belonging to LHb or MHb based on expression of *Pcdh10/Gap43* or *Tac2*, respectively. Neurons that did not express either of these markers were determined to be from surrounding areas outside of the Hb. LHb and MHb neuron clustered were filtered out for additional subclustering. The *RunUMAP()* function was used to embed neuronal nuclei into UMAP space and the *FindNeighbors()* and *FindClusters()* functions were used to cluster based on overall gene expression similarity. Known marker genes were used to assign cell type identities to each subcluster.

#### Calculation of stress DEGs within Hb clusters

To identify DEGs between control, susceptible, and resilient in LHb and MHb, we performed pseudobulk DESeq2^136^ analysis. For the LHb and MHb cluster separately, raw RNA counts were aggregated by sample ID to create pseudobulk count matrices, representing summed gene counts per sample within each cell type. The known sex chromosome markers *Xist* and *Tsix* were removed to reduce noise. For each cell type and comparison (susceptible vs control, resilient vs control, susceptible vs resilient), we created DESeq2 datasets using the pseudobulk counts and metadata. The design formula included sex as a covariate to adjust for sex-specific effects: `design = ∼ sex + condition`. Condition factors were re-leveled such that the reference group was control for comparisons to control or resilient for susceptible vs resilient. Normalized counts were calculated and genes were filtered based on expression thresholds to improve statistical power: genes with normalized counts above the 5th percentile (or a minimum of 25 counts) in at least two samples were retained. DESeq2 Wald tests were performed to identify DEGs and an adjusted p-value threshold (FDR) of 0.1 was applied to determine significance. This approach was iteratively applied to both the LHb and MHb cluster and each behavioral comparison, yielding comprehensive DEG lists.

#### Gene ontology (GO) analysis

Gene Ontology enrichment analysis was performed using the Database for Annotation, Visualization and Integrated Discovery (DAVID) to identify significantly enriched biological processes among differentially expressed genes. DEG lists were submitted using official gene symbols and analyzed against the *Mus musculus* background. Enriched GO terms were considered significant at a Benjamini-adjusted p-value (FDR) < 0.05.

#### CellChat for analysis of VTA receptor-ligand interactions

We used CellChat^106,107^ to compare inferred cell-cell interactions across the entire VTA network in each behavioral group (control, susceptible, resilient). For each group, a CellChat object was initialized using the assigned cell type identities. Overexpressed genes and ligand-receptor pairs were identified, followed by computation of cell-cell communication probabilities as previously described^106,107^. Interactions involving fewer than 10 nuclei were filtered out. The CellChat objects for each group were merged using the mergeCellChat() function for comparative visualization.

#### Identification of VTA neuronal subtypes

In the VTA dataset, clustering confirmed the presence of expected neuronal and non-neuronal populations, which we labeled based on expression of established marker genes^38,39^ (Figure 2A-B, S3A-I). These populations included neuronal and non-neuronal cell types. Clusters were identified as neuronal based on expression of *Syt1* (Figure S3A-B) and filtered out for additional subclustering. The *RunUMAP()* function was used to embed the neuronal nuclei into UMAP space and the *FindNeighbors()* and *FindClusters()* functions were used to cluster neuronal nuclei based on overall gene expression similarity. Known marker genes were used to assign identities to each subcluster (Figure 3A-E).

#### Mapping of VTA neuronal subtypes to Allen Brain Cell Atlas

To assess correspondence between experimentally derived neuronal clusters and reference-defined brain cell types, we used the Allen Institute’s Allen Brain Cell (ABC) Atlas MapMyCells tool (RRID:SCR_024672). A raw count matrix was submitted to the *mapmycells* function, which performs transcriptome-to-reference matching using a cosine similarity–based classifier trained on the ABC taxonomy. The output included the top-ranked ABC cell classes and subclasses corresponding to each input cluster, along with their relative mapping proportions. Results are summarized in Table S6, where ABC cell classes accounting for up to 90% of the mapping for each cluster are listed. This analysis was used to ensure accurate regional and neurotransmitter-level annotation.

#### Calculation of DEGs within VTA neuronal subtypes

To identify DEGs within each VTA neuronal subcluster, we performed pseudobulk DESeq2 analysis as described above.

#### LDA analysis of VTA neurons

To assess transcriptional similarity across behavioral outcomes, we performed shrinkage discriminant analysis (SDA), a regularized form of linear discriminant analysis (LDA) suited for high-dimensional data. Analyses were conducted separately for VTA DA, glutamate, and GABA6 neuron populations using normalized gene expression matrices. All SDA analyses were performed using the top 2,000 most variable genes as input features. Nuclei were grouped by outcome (control, susceptible, or resilient).

We used hold-one-sample-out cross-validation to evaluate classification performance while avoiding overfitting to individual animals. Each fold held out all nuclei from one sample, and the SDA model was trained on the remaining samples. We used the sda() function from the sda R package^137^, which implements shrinkage estimation of the within-class covariance matrix using the Ledoit-Wolf estimator.

For 2D SDA, all three groups (control, susceptible, resilient) were included in training and the top two discriminant axes (LD1 and LD2) were computed to maximize separation between all three groups. Test set nuclei were then projected into this space and assigned predicted class labels based on the trained model. Classification performance was assessed across all folds by computing confusion matrices and overall accuracy.

To examine the positioning of susceptible nuclei along the control-resilient axis, we conducted a separate 1D SDA analysis. The SDA model was trained only on control and resilient nuclei to derive a single discriminant axis (LD1). Susceptible nuclei were then projected onto this axis and their predicted labels were assigned using the trained SDA classifier.

## Notes

### Competing Interest Statement

The authors have declared no competing interest.

