## Supplemental Figures for "Stress resilience is associated with transcriptional remodeling in the VTA"

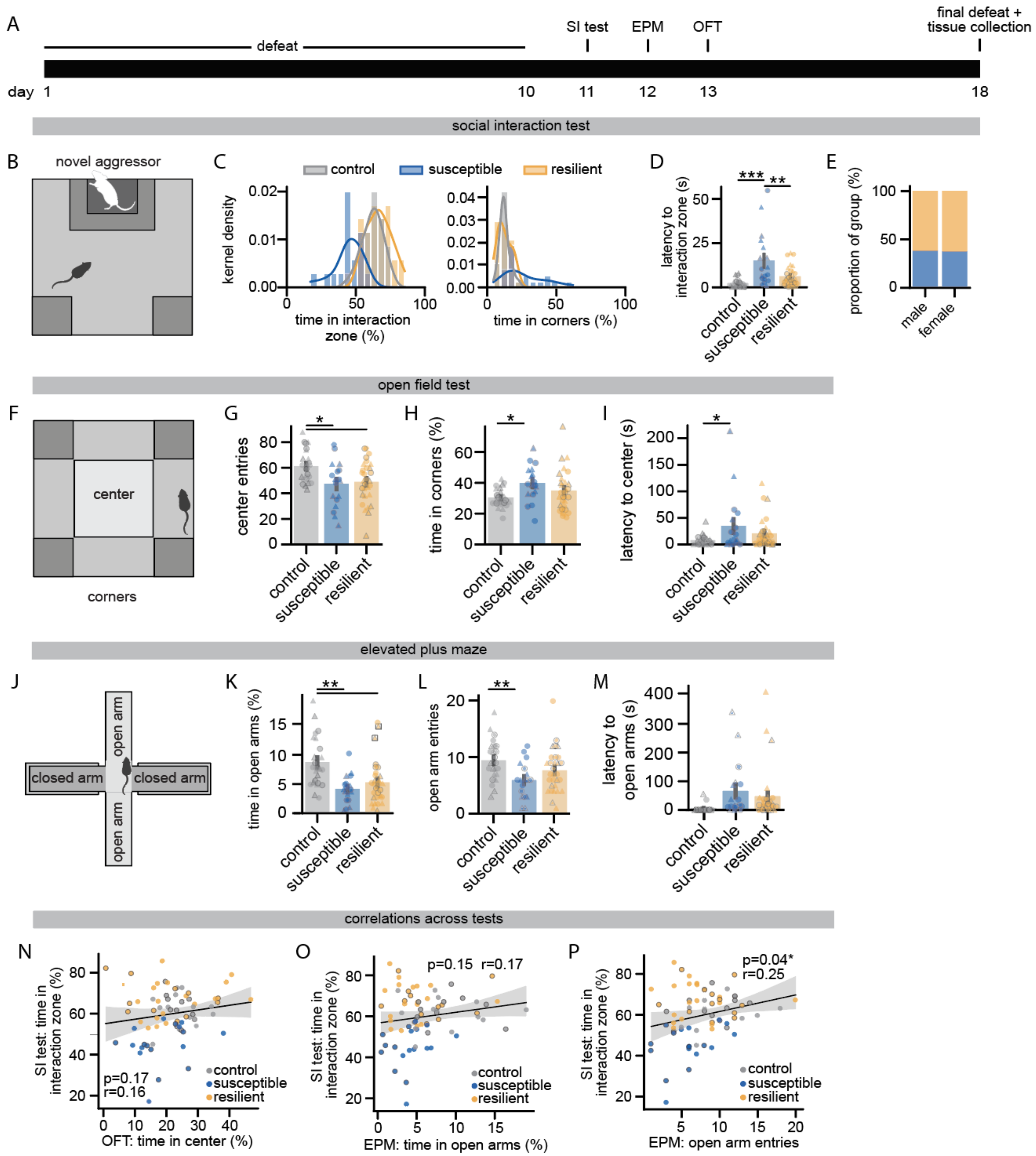


**Figure S1. Post hoc behavioral tests correlate with susceptible and resilient phenotypes from the social interaction test.**

**A.** Timeline of behavioral experiments across chronic social defeat stress. Tissue was taken ~10 minutes following a final defeat exposure.

**B.** Schematic of the social interaction (SI) test.

**C.** Distribution of time spent in the interaction zone (left) and time spent in the corners of the arena (right) during the SI test.

**D.** Latency to enter the social interaction zone during the SI test (1-way ANOVA for latency to enter interaction zone with behavior outcome as factors, $F_{(2,69)}$=11.65, $p$=4.4e-05; Tukey HSD for control vs resilient $p$=0.24, control vs susceptible $p$=0, resilient vs susceptible $p$=0.003). Black outlines indicate which mice were used for sequencing.

**E.** Proportion of each sex termed susceptible or resilient based on time spent in the interaction zone during the SI test.

**F.** Schematic of the open field test (OFT).

**G.** Number of entries into the center of the OFT (1-way ANOVA for entries into center of the OFT with behavior outcome as factors, $F_{(2,69)}$=5.60, $p$=0.006; Tukey HSD for control vs resilient $p$=0.014, control vs susceptible $p$=0.015, resilient vs susceptible $p$=0.935). Black outlines indicate which mice were used for sequencing.

**H.** Time spent in the corners of the OFT (1-way ANOVA for time in the corners of the OFT with behavior outcome as factors, $F_{(2,69)}$=3.9, $p$=0.025; Tukey HSD for control vs resilient $p$=0.278, control vs susceptible $p$=0.019, resilient vs susceptible $p$=0.298). Black outlines indicate which mice were used for sequencing.

**I.** Latency to enter the center of the OFT (1-way ANOVA for latency to enter the center of the OFT with behavior outcome as factors, $F_{(2,69)}$=3.26, $p$=0.044; Tukey HSD for control vs resilient $p$=0.356 control vs susceptible $p$=0.034, resilient vs susceptible $p$=0.348). Black outlines indicate which mice were used for sequencing.

**J.** Schematic of the elevated plus maze (EPM).

**K.** Time spent in the open arms of the EPM (1-way ANOVA for time in open arms with behavior outcome as factors, $F_{(2,69)}$=9.29, $p$=2.7e-04; Tukey HSD for control vs resilient $p$=0.003 control vs susceptible $p$=5e-4, resilient vs susceptible $p$=0.561). Black outlines indicate which mice were used for sequencing.

**L.** Number of entries into the open arms of the EPM (1-way ANOVA for number of entries with behavior outcome as factors, $F_{(2,69)}$=4.89, $p$=0.01; Tukey HSD for control vs resilient $p$=0.175 control vs susceptible $p$=0.008, resilient vs susceptible $p$=0.255). Black outlines indicate which mice were used for sequencing.

**M.** Latency to enter either of the two open arms of the EPM (1-way ANOVA for latency to enter open arms with behavior outcome as factors, $F_{(2,69)}$=3.70, $p$=0.03; Tukey HSD for control vs resilient $p$=0.107 control vs susceptible $p$=0.034, resilient vs susceptible $p$=0.712). Black outlines indicate which mice were used for sequencing.

**N.** Relationship between time in the interaction zone during the SI test and time in the center of the OFT (Pearson correlation, $r$=0.164, $p$=0.168). Black outlines indicate which mice were used for sequencing.

**O.** Relationship between time in the interaction zone during the SI test and time in the open arms of the EPM (Pearson correlation, $r$=0.170, $p$=0.153). Black outlines indicate which mice were used for sequencing.

**P.** Relationship between time in the interaction zone during the SI test and number of entries into either of the open arms of the EPM (Pearson correlation, $r$=0.247, $p$=0.036). Black outlines indicate which mice were used for sequencing.

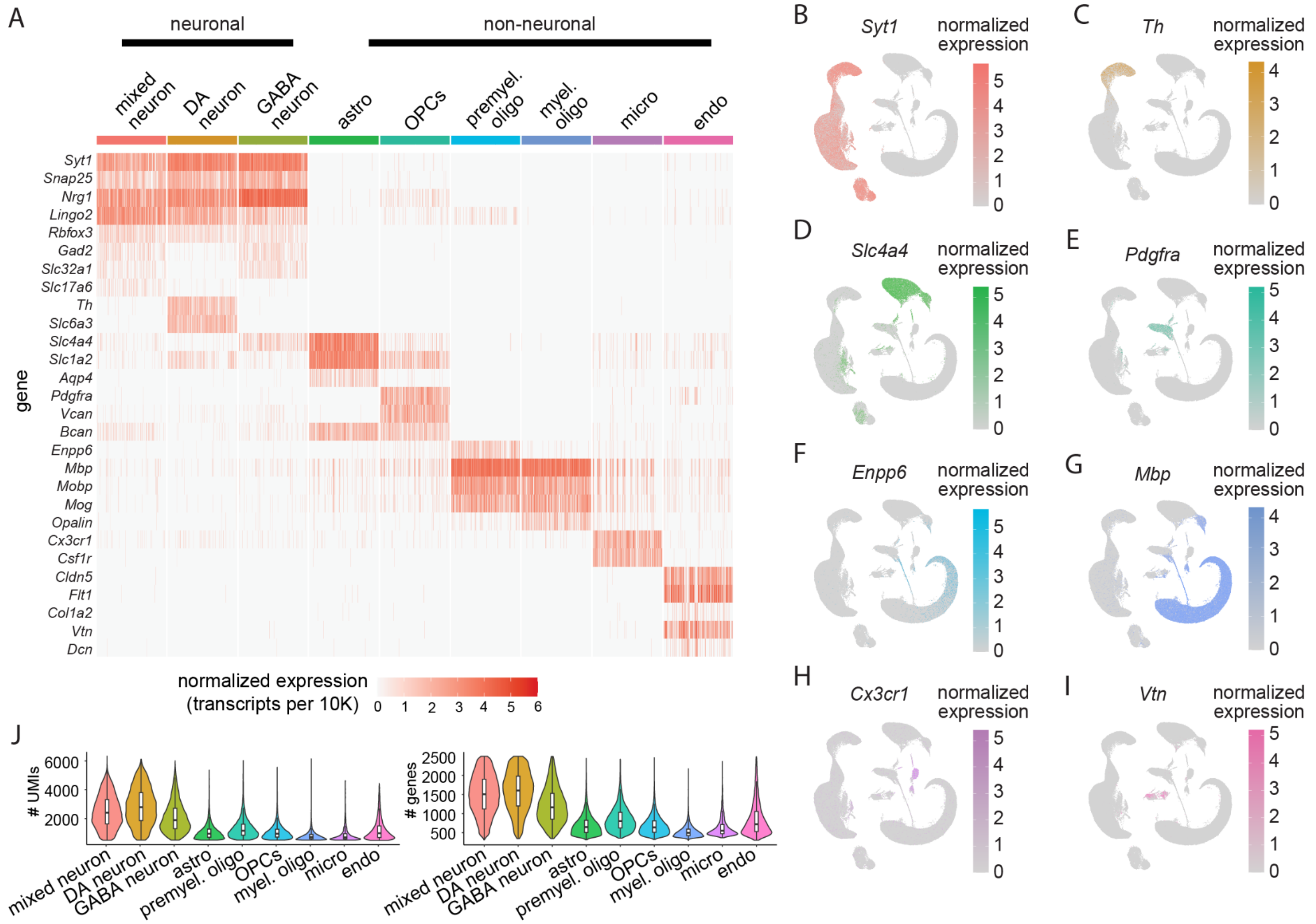


**Figure S2. Characterization and quality control metrics from snRNA-sequencing of VTA.**

**A.** Heatmap showing normalized expression of known marker genes across clusters.

**B-I.** Expression of known marker genes across UMAP space.

**J.** Violin plots showing the unique molecular identifier (UMI; top) and gene (bottom) distributions in each cell type cluster.

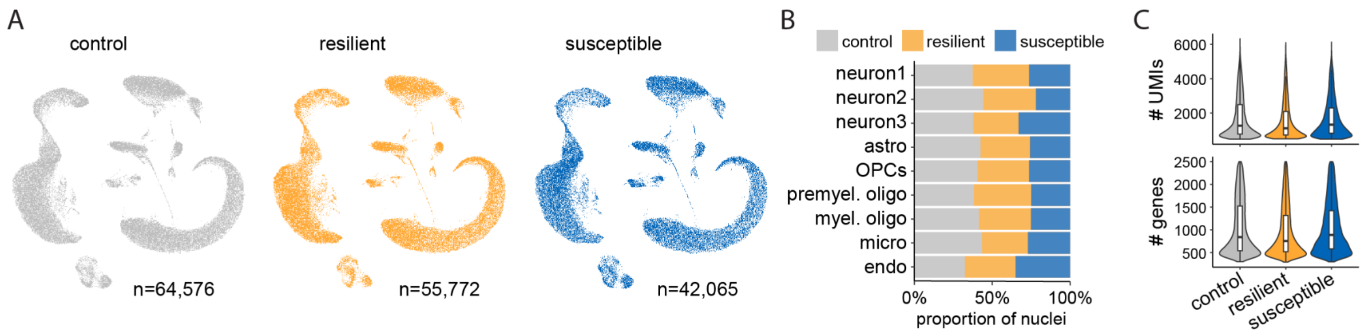


**Figure S3. Quality control metrics from snRNA-sequencing of VTA across behavioral groups.**

**A.** UMAP of all VTA nuclei with split by behavioral outcome of the sample they are from (n=64,576 control, n=55772 resilient, n=42065 susceptible).

**B.** Percentage of nuclei in each cell type from control, resilient, and susceptible samples.

**C.** Violin plots showing the UMI (top) and gene (bottom) distributions in each behavioral group.

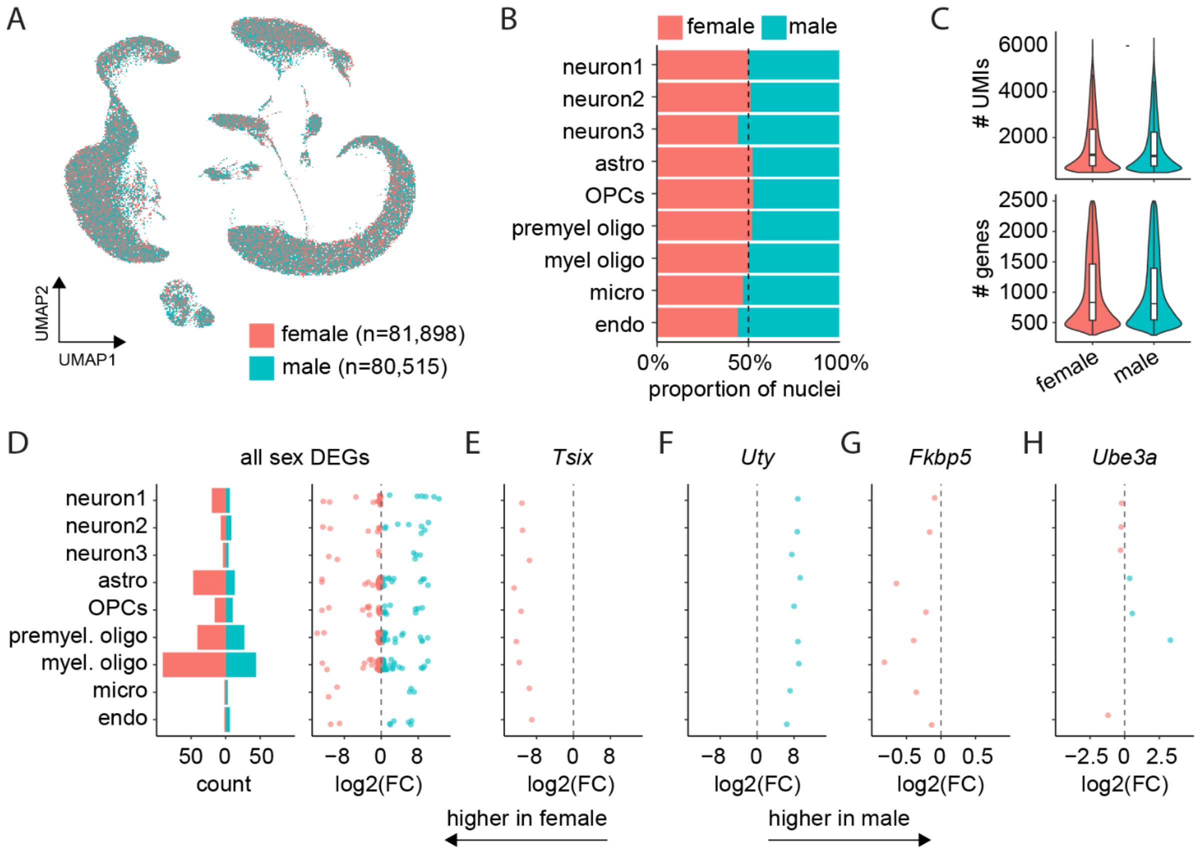


**Figure S4. Sex differences across VTA cell types.**

**A.** UMAP of all VTA nuclei with points colored by the sex of the sample they are from (n=81,898 female, n=80,515 male).

**B.** Percentage of nuclei in each cell type from male or female samples.

**C.** Violin plots showing the UMI (top) and gene (bottom) distributions in male vs female samples.

**D.** Sex DEGs by cluster. Left: number of DEGs; right: log2FoldChange of each DEGs. DEGs with a positive log2(FoldChange) are more highly expressed in males, and those with a negative log2(FoldChange) are more highly expressed in females.

**E.** Representative example of an X-linked gene known to be female biased.

**F.** Representative example of a Y-linked gene known to be male biased.

**G.** Example of a non-sex chromosome-linked gene found to be female biased.

**H.** Example of a non-sex chromosome-linked gene found to have both a sex- and cell type- dependent expression pattern.

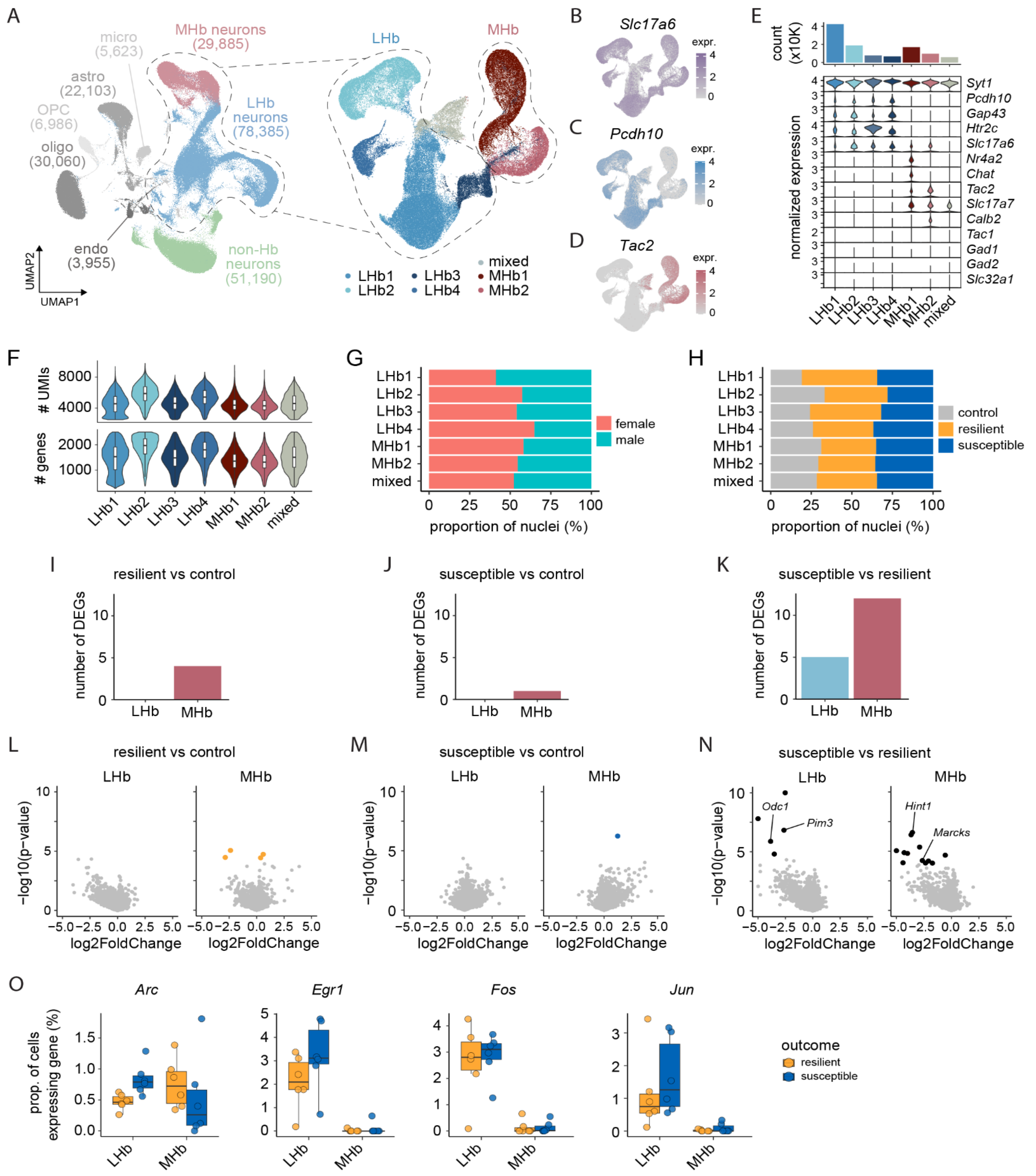


**Figure S5. Gene expression in the Hb does not uniquely define resilience or susceptibility.**

**A.** Uniform manifold approximation and projection (UMAP) of all habenula nuclei (left) and neuronal subclusters (right) with points colored by cell type classification.

**B.** Feature plot showing expression of *Slc17a6* (Vglut2) across habenula neuron UMAP space.

**C.** Same as B, but for lateral habenula marker gene *Pcdh10*.

**D.** Same as B-C, but for medial habenula marker gene *Tac2*.

**E.** Number of nuclei per habenula neuronal subcluster (top) and expression of select marker genes across habenula neuronal subclusters (bottom).

**F.** Violin plots showing the unique molecular identifier (UMI; top) and gene (bottom) distributions in each Hb neuron cluster.
**G.** Percentage of nuclei in each cluster from male or female samples.
**H.** Percentage of nuclei in each cluster from control, resilient, and susceptible samples.
**I.** Barplot showing the number of DEGs in the main LHb and MHb clusters from panel A (left) in resilience compared to control (LHb=0, MHb=4).
**J.** Barplot showing the number of DEGs in the main LHb and MHb clusters from panel A (left) in susceptibility compared to control (LHb=0, MHb=1).
**K.** Barplot showing the number of DEGs in the main LHb and MHb clusters from panel A (left) in susceptibility compared to resilient (LHb=5, MHb=12).
**L.** Volcano plots showing DEGs between resilient and control in LHb (left) and MHb (right). Nominal p-values are plotted, but points are colored if the adjusted p-value (Bonferroni post hoc correction) is <0.1.
**M.** Same as L, but for DEGs between susceptible and control.
**N.** Same as L-M, but for DEGs between susceptible and resilient.
**O.** Percent of nuclei per sample in LHb and MHb expressing each IEG.

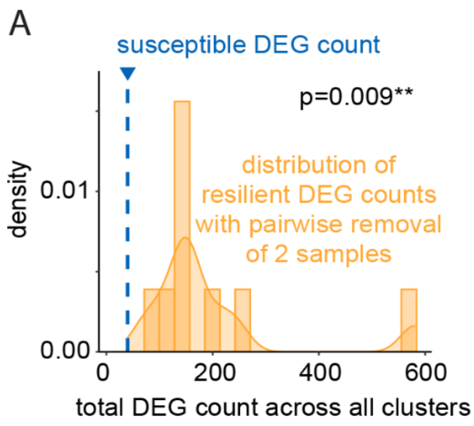


**Figure S6. Differences in resilient vs control and susceptible vs control DEG counts is not due to sample size differences.**

**A.** Distribution of total DEG counts across all cell types in resilient vs control comparisons after removing each combination of one male and one female sample. Dashed line indicates the DEG count for susceptible vs control. The number of susceptible vs control DEGs is significantly less than the distribution of resilient vs control DEG counts (one-sample Wilcoxon signed-rank, $p$=0.009).

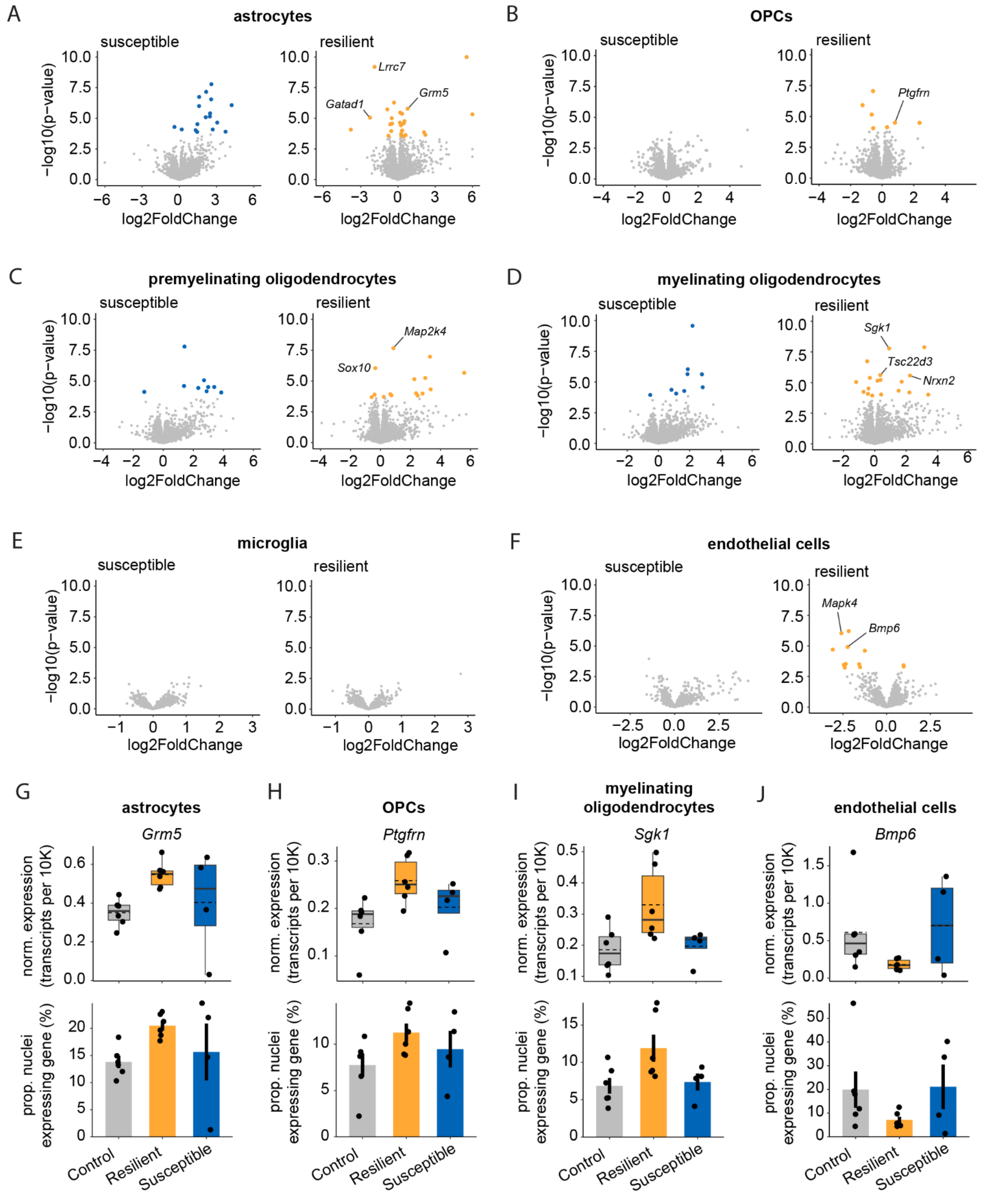


**Figure S7. Example DEGs in non-neuronal cell types of the VTA.**

**A.** Volcano plots showing differentially expressed genes in astrocytes in susceptible vs control (left) and resilient vs control (right). Nominal p-values are plotted, but points are colored if the adjusted p-value (Bonferroni post hoc correction) is <0.1.

**B-F.** Same as A, but for OPCs (B), premyelinating oligodendrocytes (C), myelinating oligodendrocytes (D), microglia (E), and endothelial nuclei (F).

**G.** Example astrocyte DEG *Grm5*; top: normalized expression across samples; bottom: percentage of astrocyte nuclei expressing *Grm5* per sample.
**H.** Same as G, but for *Ptgfrn* in OPCs.
**I.** Same as G, but for *Sgk1* in myelinating oligodendrocytes.
**J.** Same as G, but for *Bmp6* in endothelial nuclei.

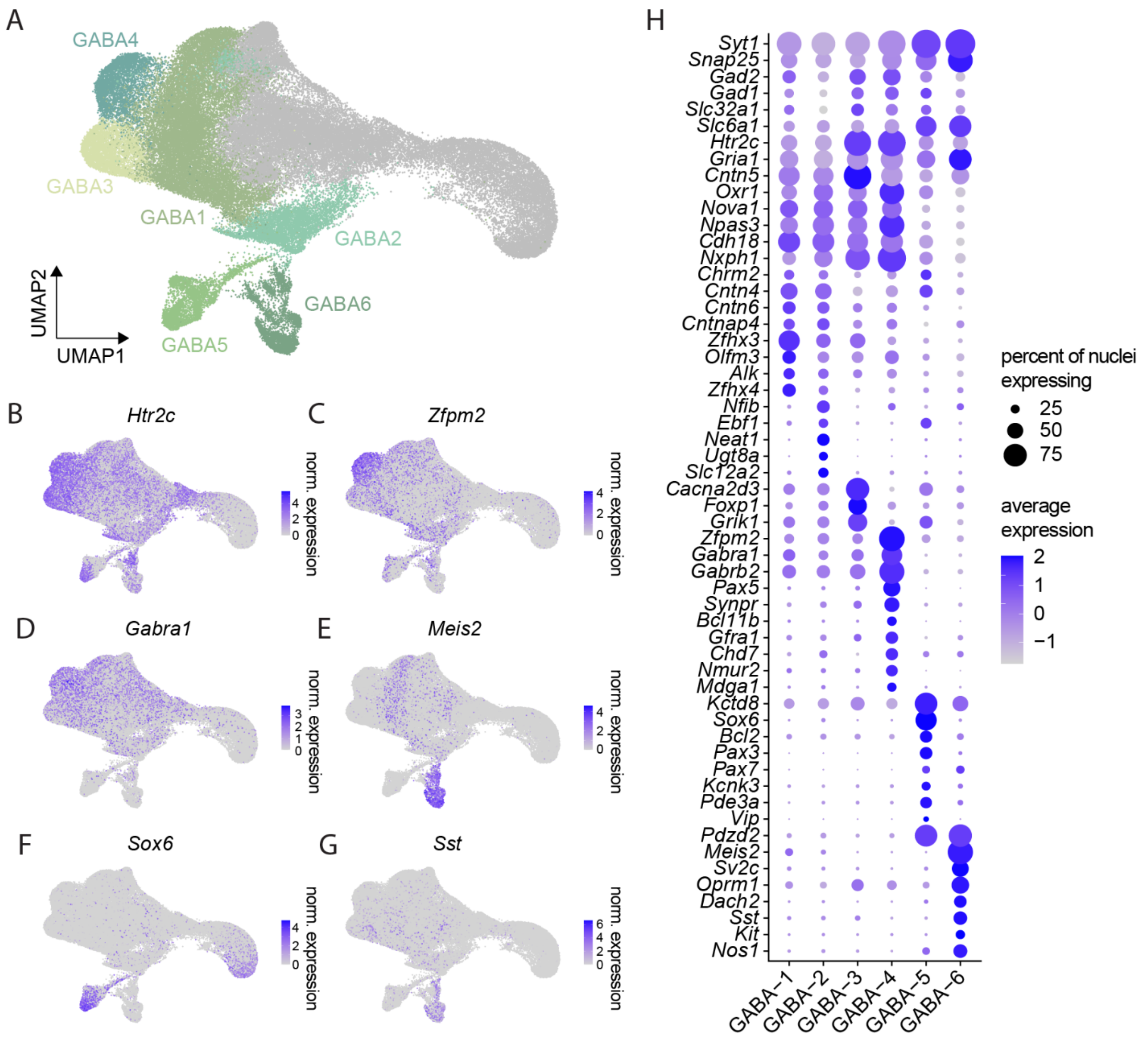


**Figure S8. GABAergic neuron diversity in the VTA.**

**A.** UMAP of VTA neuron nuclei with GABA subclusters highlighted.

**B-G.** Distribution of GABA marker genes in UMAP space.

**H.** Dot plot showing the expression of traditional GABAergic and additional GABA subcluster marker genes. Size of dot represents the proportion of nuclei in each cluster expressing the gene and transparency represents the average expression of the gene in each cluster.

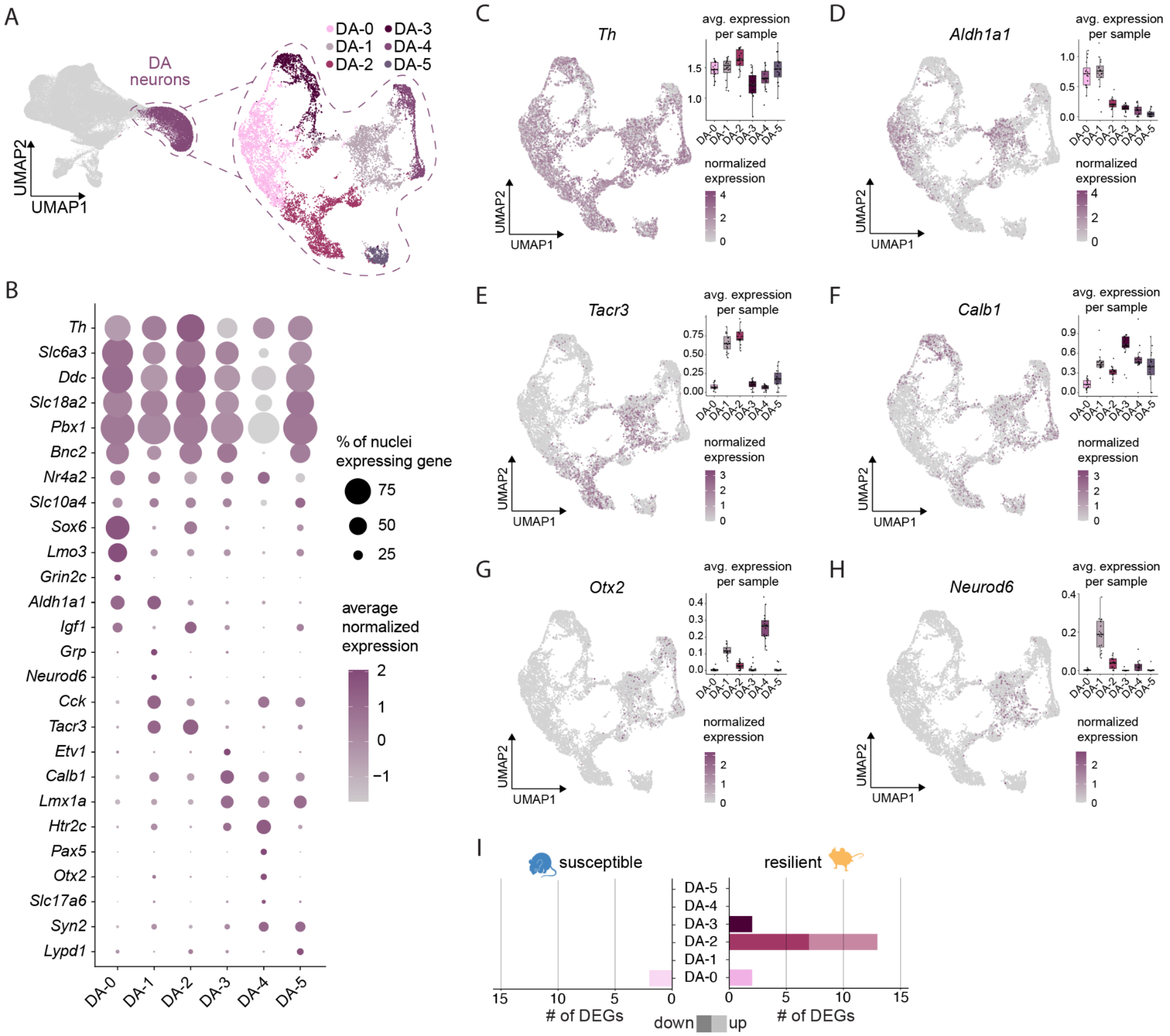


**Figure S9. DA neuron diversity in the VTA.**

**A.** UMAP of VTA DA neurons sub-clustered into dopaminergic subtypes, with points colored by subcluster.

**B.** Dot plot showing the expression of traditional dopaminergic and additional DA subcluster marker genes. Size of dot represents the proportion of nuclei in each cluster expressing the gene and transparency represents the average expression of the gene in each cluster.

**C-H.** Distribution of select DA marker genes in UMAP space, with insets showing average normalized expression of each gene across subclusters.

**I.** Barplot showing the number of DEGs in each DA subcluster compared to control in susceptible (left) and resilient (right).
